# Inference of recombination maps from a single pair of genomes and its application to archaic samples

**DOI:** 10.1101/452268

**Authors:** Gustavo V. Barroso, Natasa Puzovic, Julien Y. Dutheil

## Abstract

Understanding the causes and consequences of recombination rate evolution is a fundamental goal in genetics that requires recombination maps from across the tree of life. Since statistical inference of recombination maps typically depends on large samples, reaching out studies to non-model organisms requires alternative tools. Here we extend the sequentially Markovian coalescent model to jointly infer demography and the variation in recombination along a pair of genomes. Using extensive simulations and sequence data from humans, fruit-flies and a fungal pathogen, we demonstrate that iSMC accurately infers recombination maps under a wide range of scenarios – remarkably, even from a single pair of unphased genomes. We exploit this possibility and reconstruct the recombination maps of archaic hominids. We report that the evolution of the recombination landscape follows the established phylogeny of Neandertals, Denisovans and modern human populations, as expected if the genomic distribution of crossovers in hominids is largely neutral.

---

Meiotic recombination is a major driver of the evolution of sexually-reproducing species^1^. The crossing-over of homologous chromosomes creates new haplotypes and breaks down linkage between neighbouring loci, thereby impacting natural selection^2,3^ and consequently the genome-wide distribution of diversity^4^. The distribution of such cross-over events is heterogeneous within and among chromosomes^5,6^, and commonly referred to as the recombination landscape – apicture of how often genetic variation is shuffled in different parts of the genome. Interestingly, this picture is not static, but instead is an evolving trait that varies between populations^7,8^ and species^9^. Moreover, the proximate mechanisms responsible for shaping the recombination landscape vary among *taxa*. For example, among primates (where the PRDM9 gene is a key player determining the location of so-called recombination hotspots^10^) the landscape is conserved at the mega-base (Mb) scale, but not at the kilo-base (kb) scale^11^. In birds, which lack PRDM9, the hotspots are found near transcription start sites in the species that have been studied so far^8,12^. In *Drosophila* (where clear hotspots appear to be absent^13^), inter-specific changes are associated with mei-218 variants^14^, a gene involved in the positioning of double-strand breaks^15^. The molecular machinery influencing the distribution of cross-over events is still poorly understood in many other groups, where a picture of the recombination landscape in closely related species is lacking.

Aside from their intrinsic value in genetics, accurate recombination maps are needed to interpret the distribution of diversity along the genome. Since the rate of recombination determines the extent to which linked loci share a common evolutionary history^16^, inferring selection^17–19^, introgression^18,20^ and identifying causal loci in association studies requires knowledge of the degree of linkage between sites^21^. Furthermore, recombination can cause GC-biased gene conversion ^22,23^, which can mimic the effect of selection^24^ or interfere with it^25^. Obtaining recombination maps, however, remains a challenging task. Due to the typically low density of markers, experimental approaches provide broad-scale estimates and are limited in the number of amenable *taxa*. Conversely, population genomic approaches based on coalescent theory^26,27^ have proved instrumental in inferring recombination rates from polymorphism data.

Traditionally, population genomic methods infer recombination maps from variation in linkage disequilibrium (LD) between pairs of single nucleotide polymorphisms (SNPs)^28–30^. However, since “LD-based” methods typically require large sample sizes per population (from a dozen individuals^31^), their application is restricted to a few model organisms where such sequencing effort could be afforded. Moreover, because computation of the exact likelihood is intractable for long sequences, the underlying model only accounts for pairs of SNPs^32^, multiple pairs being considered independent and combined using a composite likelihood approach. To avoid over-fitting due to the high number of parameters, smoothness in the landscape is enforced by using heuristic corrections that have been fine-tuned for humans^28^ or fruit-flies^30^. Application of these methods to systems with distinct genomic architectures therefore requires extra-benchmarking (see for instance Stukenbrock and Dutheil^31^).

Here we introduce a new modeling framework (iSMC) to infer the variation in the recombination rate along the genome, using a single pair of unphased genomes. Using simulations, we show that iSMC is able to accurately recover the recombination landscape under diverse scenarios. We further demonstrate its efficacy with case studies in Humans, Fruit-flies and the fungal pathogen *Zymoseptoria tritici*, where experimental genetic maps are available. Finally, we exploit our new method to investigate the recombination landscape of archaic hominid samples: Ust’Ishim, the Vindija Neandertal, the Altai Neandertal and the Denisovan. Because it allows inference from datasets for which sample size is intrinsically limited, such as ancient DNA samples, our method opens a new window in the study of the recombination landscape evolution.

## RESULTS

### Overview of iSMC

Besides its common interpretation as a backwards-in-time process, the coalescent with recombination^33,34^ can also be modelled as unfolding spatially along chromosomes^35^. Starting from a genealogy at the first position of the alignment, the process moves along the chromosome sequence, adding recombination and coalescence events to the ensuing ancestral recombination graph (ARG)^36,37^ (Figure 1a). Due to long-range correlations imposed by rare recombination events that happen outside the ancestry of the sample (in so-called trapped non-ancestral material^38^), the genealogy after a recombination event cannot be entirely deduced from the genealogy before, rendering the process non-Markovian. The sequentially Markovian coalescent process (SMC)^39,40^ ignores such recombination events, but captures most of the properties of the original coalescent^41^ while being computationally tractable. It is the foundation of recent models for demographic inference^42–44^ and has been used to infer the broad-scale recombination map of the human-chimpanzee ancestor based on patterns of incomplete lineage sorting^45,46^. In the SMC, transition probabilities between genealogies are functions of ancestral coalescence rates and – of key relevance to this study – the population recombination rate (ρ = 4.Ne.r)^43,44^. Thus, heterogeneous recombination landscapes affect the SMC by modulating the frequency of genealogy transitions: genomic regions with higher recombination rate are expected to harbour relatively more genealogies than regions with smaller recombination rate (Figure 1). We leverage this information by extending the SMC to accommodate spatial heterogeneity in ρ (see Methods). In brief, our new model combines the discretised distribution of times to the most recent common ancestor (TMRCAs) of the pairwise SMC^43^ with a discretised distribution of ρ to jointly model their variation along the genome. Since we model the transition between discretised ρ categories as a spatially Markovian process along the genome, combining the SMC with the Markov model of recombination variation leads to a Markov-modulated Markov model. We cast it as a hidden Markov model^47,48^ (HMM) to generate a likelihood function, where the observed states are orthologous nucleotides and the hidden states are {TMRCA, ρ-category} pairs (Figure 1c). We name our new approach “integrative sequentially Markov coalescent (iSMC)”, as it enables jointly capturing the effect of time and space in the Coalescent. This framework explicitly connects the genealogical process with the classical definition of LD as the non-random association of alleles at different loci^49^, which has been formulated in terms of covariances in coalescence times^50^. Henceforth, we restrict the use of the term LD to its “topological” interpretation^51^.

**Figure 1.**
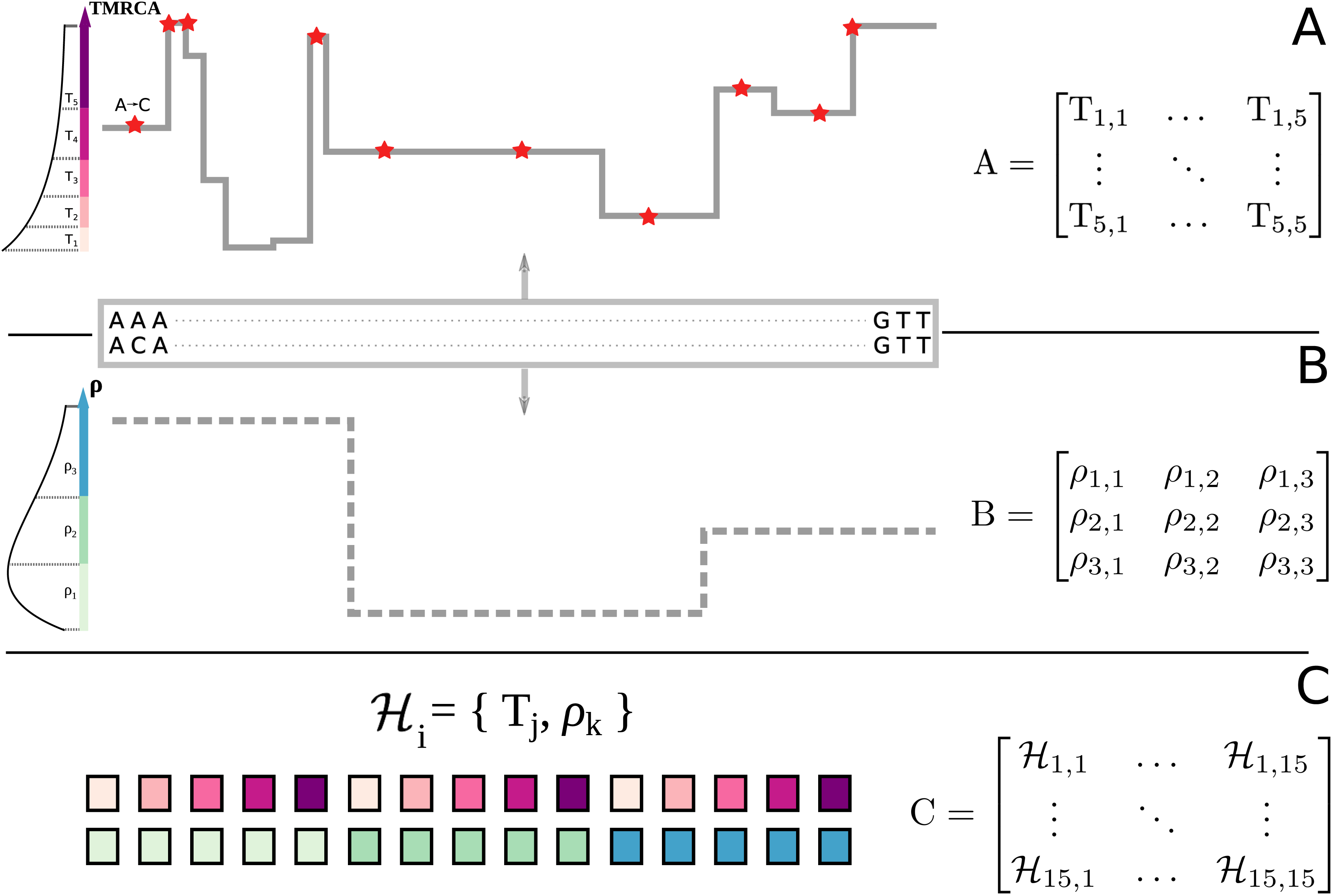
**Schematic representation of iSMC for one pair of genomes, with five time intervals and three recombination rate categories. A**, In the SMC process, the spatial distribution of TMRCAs can be described by a matrix of transition probabilities that depend on local population recombination rate ρ and the ancestral coalescence rates. **B**, variation in ρ along the genome, modelled as a Markovian process and described by a matrix of transition probabilities. **C**, the combination of both Markovian processes leads to a Markov-modulated Markovian process. The hidden states of the resulting hidden Markov model are all pairwise combinations of discretized classes in **A** and **B**.

iSMC models spatial variation in the recombination rate using a single discrete distribution (Figure 1b), which can be accommodated to various models of recombination rate variation (see Methods). After fitting the alternative distributions to sequence data, Akaike’s Information Criterium (AIC)^52^ is employed as a mean of model selection. If AIC favours a spatially heterogeneous model over the null model where ρ is constant along the genome, iSMC then estimates a recombination landscape of single-nucleotide resolution by weighting the discretised values of the selected distribution with their local posterior probabilities. In the following section, we benchmark our model on different simulated scenarios. The correlations between simulated and inferred maps reported therein were computed after binning the inferred landscapes into windows of 50 kb, 200 kb, 500 kb and 1 Mb.

### Simulation study

To assess iSMC’s overall performance, we used SCRM^53^ to simulate five recombination landscapes corresponding to different patterns of magnitude and frequency of change in ρ and a “null” scenario with constant recombination rate along the genome (see Methods). For each landscape, we simulated 10 ARGs, each describing the ancestry of 2 haploid chromosomes. We tested two discretisation schemes for the joint distribution of TMRCAs and recombination rates: the first with 40 time intervals, five ρ categories; the second with 20 time intervals, 10 ρ categories, leading to a total of 200 hidden states in both configurations (**Supplemental Note**). Model selection based on AIC favours the correct model in 45 of the 50 datasets (**Table S1**), with the five exceptions belonging to the scenario where changes are frequent and of small magnitude. In this regime, transitions to regions of slightly different recombination rates do not significantly skew the distribution of genealogies, and the short length of blocks with constant ρ leaves little signal in the data. Nevertheless, correlations between simulated and inferred maps are highly significant (ranging from 0.385 to 0.563 in the five identifiable replicates with frequent changes of low magnitude, and from 0.682 to 0.928 in the others, **Tables S2, S3**). Overall, the results are consistent between replicates and robust to the choice of discretisation (Figure 2b). We use the configuration with 40 time intervals and five ρ categories, as it implements a finer discretisation of time that is more adequate to capture the effect of ancestral demography. In the following, as we introduce new simulated scenarios, we focus on the recombination landscape with frequent changes of large magnitude.

**Figure 2.**
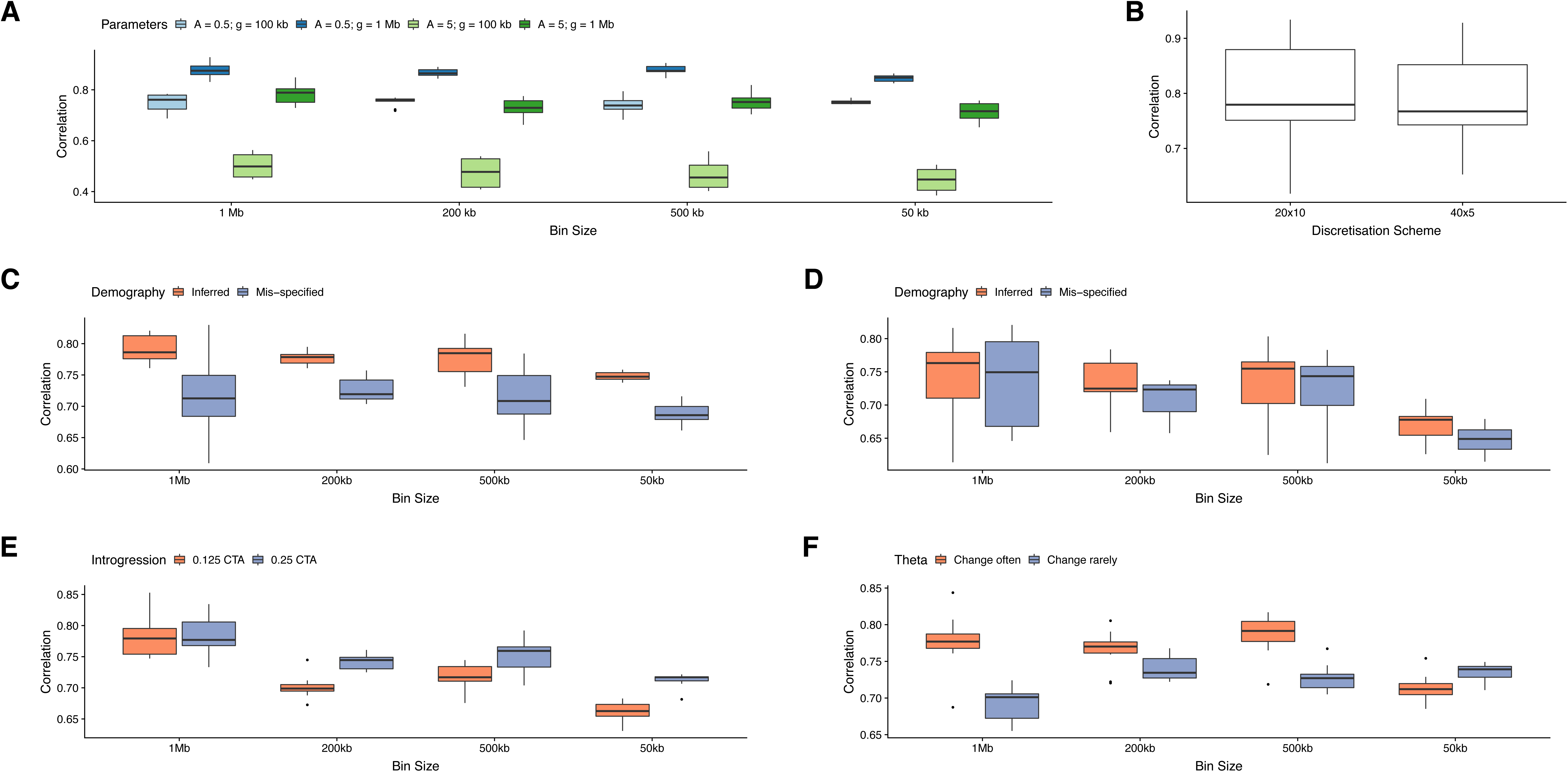
**Distribution of correlation coefficients in the simulation study.** Center line, median; box limits, upper and lower quartiles; whiskers, 1.5x interquartile range; points, outliers. **A**, four scenarios of spatial variation in the recombination rate, corresponding to different combinations of parameters (colour). **B**, the respective comparison between two discretisation schemes in the four recombination landscapes. **C-D**, comparison between a model where demography is mis-specified and another where it is jointly inferred (colour), in scenarios of recent growth (**C**) or ancient bottleneck (**D**). **E**, two scenarios of introgression, varying the time of gene-flow (colour). **F**, two scenarios of spatial variation in the mutation rate, varying its frequency of change (colour).

*Demographic history*. The random sampling of haplotypes during population bottlenecks and expansions affects LD between SNPs, thus creating spurious signals of variation in ρ^54–56^. To test whether iSMC could capture the effect of demography on the inference of recombination maps, we simulated a heterogeneous recombination landscape coupled with either a recent 20-fold increase, or ancient 20-fold decrease in population size. We then fit our model twice for each scenario: first, erroneously assuming a flat demographic history; second, allowing iSMC to infer piecewise constant coalescence rates in order to accommodate population size changes. The correlation coefficients are all highly significant (Figure 3), showing that the inferred recombination landscape is relatively robust to misspecification of the demographic scenario, but are systematically higher when demography is jointly inferred (Figure 2c-d, **Table S4, S5**). The difference is stronger at the fine scale, where, in the presence of complex demography the distribution of genealogies can get locally confined to a time period, and ignorance about differential coalescence rates reflects poorly on local ρ estimates. We conclude that the joint-inference approach of iSMC can disentangle the signal that variable recombination and fluctuating population sizes leave on the distribution of SNPs.

**Figure 3.**
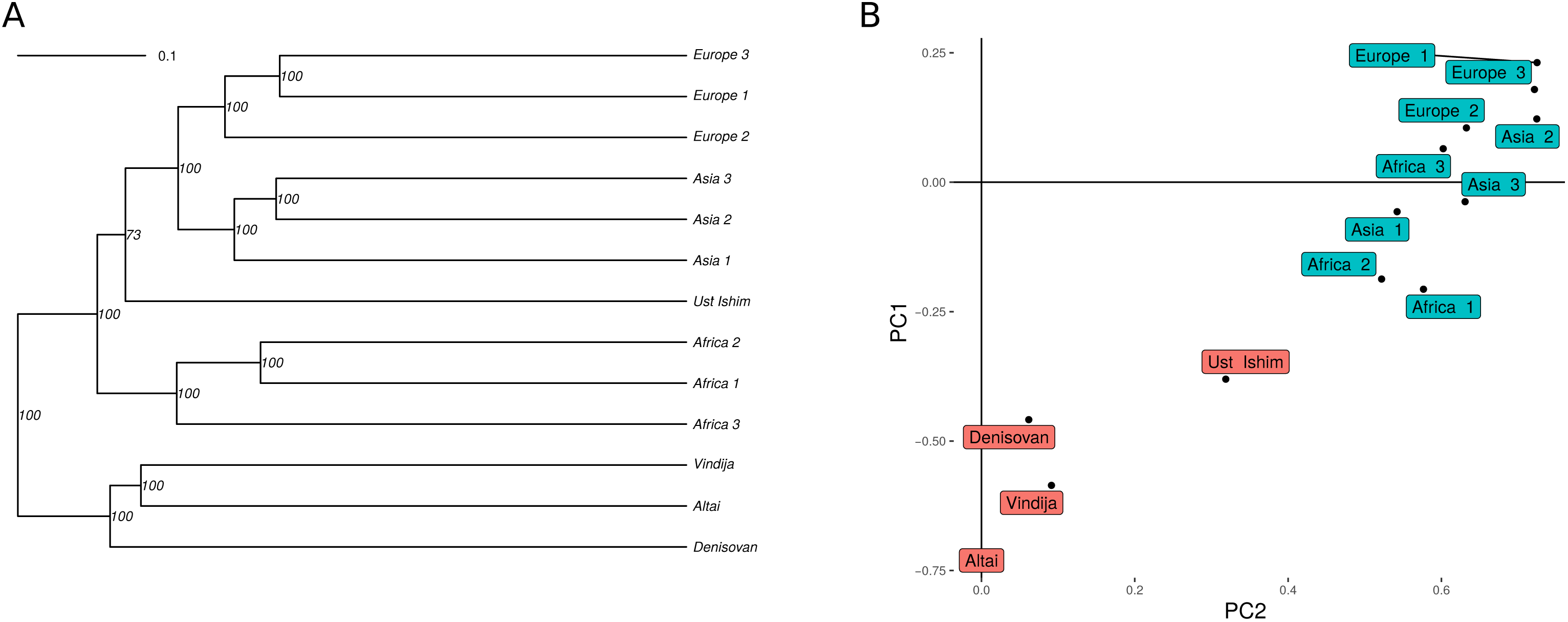
**Inference of recombination maps in the presence of recent population growth.** Each column represents a bin size (50 kb, 200 kb, 500 kb and 1 Mb). **A-D**, Simulated (black) and inferred (orange) recombination maps. **E-H**, scatter-plots of the same data. Red lines represent y = x, blue lines represent loess smoothing curve.

*Introgression events.* Recent studies suggested that introgression is a frequent phenomenon in nature^57,58^. The influx of a subset of chromosomes from a source population into another, target population (in a process analogous to a genetic bottleneck) introduces long stretches of SNPs in strong LD. Past introgression events will thus affect runs of homozygosity, potentially biasing the distribution of genealogies. To test the robustness of iSMC to the confounding effect of introgression, we simulated two scenarios of admixture which differ in their time of secondary contact between populations. The correlations between simulated and inferred recombination maps remain highly significant (ranging from 0.631 to 0.834, Figure 2e, **Table S6, S7**) and depend on the time when introgression occurred. Recombination maps are less accurately recovered in case of recent introgression, because in such case there has been less time for recombination to break SNP associations that do not reflect local ρ in the target, sampled population.

*Variation in mutation rate.* The rate of *de novo* mutations varies along the genome of many species. For example, CpG di-nucleotides experience an increase in mutation rate (μ) as a result of methylation followed by deamination into thymine, whereas the efficiency of the molecular repair machinery is negatively correlated with the distance from the DNA replication origin, causing μ to vary accordingly^59^. Such heterogeneity could bias iSMC’s estimates because the transition into a region of higher μ mimics the transition to a genealogy with a more ancient common ancestor, since in both cases the outcome is locally increased genetic diversity. To assess the impact of variation of mutation rate on the estimation of recombination rate, we simulated two scenarios of variation of θ = 4.Ne.μ along the genome, corresponding to low and high frequency of change, relative to the frequency of change in the recombination rate. We report that transitions to different mutation rates along the genome globally do not introduce substantial biases in our estimates (correlations range from 0.655 to 0.844, Figure 2f, **Table S8, S9**). There is, however, an effect of the local ratio θ / ρ at the fine-scale: regions with a ratio < 1 tend to have their local ρ underestimated because the relatively small number of mutations render some recombination events undetectable (**Figure S1a**). Importantly, this effect is a consequence of the magnitude of μ, and is not expected as a result of the local reduction in diversity caused by selection, since in the latter case there is a linear scaling of both θ and ρ (due to a decrease in the local effective population size, Ne), keeping their ratio constant. Furthermore, the bias is reduced when site-specific recombination estimates are binned into larger windows (**Figure S1b**).

**Application to a fungal pathogen, fruit-flies and humans**

We benchmarked iSMC on model organisms with contrasting genomic architectures and evolutionary histories. The leaf blotch *Zymoseptoria tritici* is a highly polymorphic fungal pathogen with a compact genome (40 Mb) that is under widespread selection^60^ and exhibits an extremely rugged recombination landscape^31,61^. In this species, AIC favours a heterogeneous model with the presence of recombination hotspots in all three pairs of genomes analysed (Table 1, see Methods). Under this model, correlations between iSMC maps and a previously published genetic map^61^ are highly significant (**Table S10**) at the 20 kb scale (0.279, 0.287 and 0.308) and increase at the 100 kb scale (0.393, 0.476 and 0.499). In sharp contrast to *Z. tritici*, the recombination landscape in *Drosophila* is notably smooth^13^, and AIC favours a heterogeneous model based on a Gamma distribution. Under this model, correlations between iSMC maps and a genetic map of this species are highly significant (**Table S10**) at the 100 kb scale (0.334, 0.342 and 0.363), and increase at the 1 Mb scale (0.549, 0.613 and 0.644). Like in *Zymoseptoria tritici*, model fitting in humans favours a heterogeneous distribution of recombination rates with the presence of hotspots (Table 1). We inferred recombination maps under this model for each of the three Yoruban (African), three Dai Chinese (Asian) and three Finnish (European) genomes available in the Simons Genome Diversity Project^62^, and correlated them to the sex-averaged genetic map from DECODE^7^, after binning into windows of 50 kb, 200 kb, 500 kb, and 1 Mb (Table 2, **Table S11**). Correlations between the DECODE map and iSMC maps from the African individuals(range from 0.116 to 0.388 depending on window size) are lower than when individual maps from either Asia (0.177 to 0.457) or Europe (0.179 to 0.455) are compared. This was expected since the DECODE map was estimated from a pedigree study of the Icelandic population. The correlations increase if we use consensus maps (defined here as the average maps among three diploids) from each population (from 0.177 to 0.431 according to window size for Africans, 0.251 to 0.508 for Asians and 0.253 to 0.521 for Europeans). Taken together these results show that iSMC can infer recombination maps from species with extremely different recombination profiles.

**Table 1:** **Performance of iSMC in three distantly related species.** AIC and Kendall correlations between individual recombination maps and a genetic map from each species.

|  |  | **Drosophila** (chr 2L)<br>100-kb / 1-Mb |  |  | **Zymoseptoria** (chr 1)<br>20-kb / 200-kb |  |  | **Humans** (chr 10)<br>50-kb / 1-Mb |  |  |
| --- | --- | --- | --- | --- | --- | --- | --- | --- | --- | --- |
|  | Configuration | Sample 1 | Sample 2 | Sample 3 | Sample 1 | Sample 2 | Sample 3 | Sample 1 | Sample 2 | Sample 3 |
| **AIC** | 40x5 | **2240174.6** | **2247931.8** | **2207295.4** | 572264.6 | 589366.4 | 589410.5 | 1399031.9 | 1374656.2 | 1434109.5 |
|  | 40x(4+1) | 2240269.1 | 2247937.7 | 2207417.0 | 572101.7 | 589772.2 | 589614.0 | 1399099.2 | 1374692.4 | 1434148.8 |
|  | 40x2 | 2240267.8 | 2248028.2 | 2207415.9 | 571974.0 | 589392.2 | 589509.4 | 1399096.3 | 1374690.3 | 1434146.9 |
|  | 100x2 | 2240359.0 | 2247948.0 | 2207537.0 | **571756.1** | **588792.1** | **588748.8** | **1398867.6** | **1374403.8** | **1433950.9** |
| **Cor.**<br>**fine-scale** | 40x5 | **0.363\*\*\*** | **0.334\*\*\*** | **0.342\*\*\*** | 0.041 | 0.118** | 0.245*** | 0.189*** | 0.196*** | 0.192*** |
|  | 40x(4+1) | 0.294*** | 0.267*** | 0.300*** | 0.216*** | 0.272*** | 0.282*** | 0.196*** | 0.191*** | 0.179*** |
|  | 40x2 | 0.293*** | 0.261*** | 0.300 | 0.217*** | 0.225*** | 0.223*** | 0.198*** | 0.192*** | 0.180*** |
|  | 100x2 | 0.289*** | 0.265*** | 0.298*** | **0.287\*\*\*** | **0.279\*\*\*** | **0.308\*\*\*** | **0.211\*\*\*** | **0.190\*\*\*** | **0.179\*\*\*** |
| **Cor.**<br>**large-scale** | 40x5 | **0.613\*\*\*** | **0.549\*\*\*** | **0.644\*\*\*** | -0.306* | -0.007 | 0.191 | 0.445*** | 0.436*** | 0.460*** |
|  | 40x(4+1) | 0.502*** | 0.455** | 0.494*** | 0.421*** | 0.471*** | 0.499*** | 0.449*** | 0.445*** | 0.452*** |
|  | 40x2 | 0.510*** | 0.423** | 0.494*** | 0.398** | 0.476*** | 0.421*** | 0.448*** | 0.449*** | 0.450*** |
|  | 100x2 | 0.494*** | 0.439** | 0.486*** | **0.499\*\*\*** | **0.393\*\*** | **0.476\*\*\*** | **0.455\*\*\*** | **0.445\*\*\*** | **0.446\*\*\*** |

\* p-value < 0.05; \*\* p-value < 0.01; \*\*\* p-value < 0.001

**Table 2:**
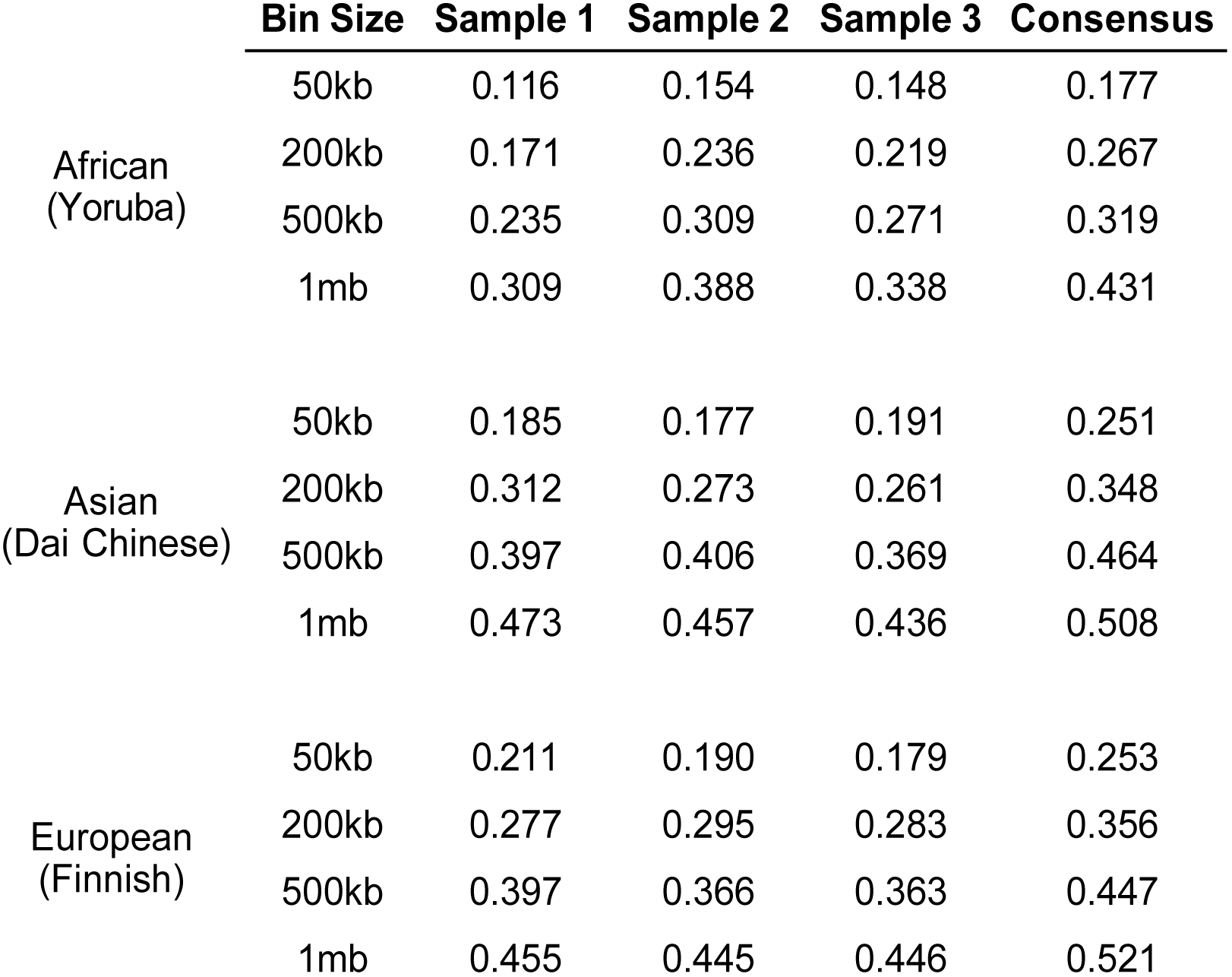
**Performance of iSMC in distinct human populations.** Kendall correlations between individually inferred recombination maps for each of three human populations and the DECODE genetic map. All values are significant with p-value < 0.001.

### Application to archaic samples

At the fine scale, the location of cross-over events in great apes is strongly influenced by the sequence of the PRDM9 gene^6,10,63^. Recombination hotspots determined by PRDM9 tend to erode over time, being replaced somewhere else in the genome with the rise of new PRDM9 alleles. Such turn-over of hotspot locations leads to rapid evolution of the recombination landscape^64,65^, therefore, if the location of hotspots is selectively neutral, recombination maps should become more dissimilar with increasing divergence between populations and species. This hypothesis has been corroborated by two lines of evidence. First, comparisons between recombination maps of extant great ape species show no overlap of hotspots at the fine scale, but correlations increase with window size. While this observation suggests that molecular players other than PRDM9 shape the landscape at the large scale, such positive relationship could be an artefact of reduced variance when recombination estimates are averaged in larger windows (e.g., correlations increase with window size in our simulations, despite recombination landscape being static over time, **Table S1-2**). Second, *in silico* prediction of PRDM9 binding sites in the Denisovan genome has shown no overlap of hotspots with modern humans^66^. iSMC’s unique ability to extract information from single diploids allowed for an alternative test of this hypothesis through the analyses of four archaic samples^67^: the Altai Neandertal^68^, the Vindija Neandertal^69^, the Denisovan^70^ and the Ust’Ishim individual^71^, a 45,000-year-old modern human from Siberia. Pairwise correlations between individual maps at the fine (50 kb) scale reveals that the evolution of the recombination landscape recapitulates the evolutionary history of hominids: Asians and Europeans form a distinct cluster; the 45,000 year-old Ust’Ishim is a long branch, sister to this clade, depicting similarities in the recombination landscape that have been frozen by his demise soon after the out-of-Africa migration; and all modern humans are highly diverged from the monophyletic Neandertal-Denisovan group (Figure 4a). The tree has high bootstrap support for its branches and is corroborated by principal component analysis (Figure 4b). Overall, the topology is consistent at larger scales (**Figure S2**), with the notable exception of Ust’Ishim forming a highly supported cluster with the Altai Neandertal. Interestingly, introgression from Neandertals is estimated to have occurred into the population ancestral to this ancient *Homo sapiens*, 232-430 generations before his lifetime. His DNA carries segments of Neandertal ancestry that are longer and more abundant than those in contemporary modern humans^71^. It is thus tempting to interpret this result as a consequence of the hybridisation event. Besides similar LD in the longer introgressed regions, a non-exclusive possibility is that the recombination landscape of Ust’Ishim was influenced by trans-acting alleles inherited from Neandertals, presumably acting at a larger scale than PRDM9. This could explain the discrepancy between recombination-based trees at the large *versus* fine scale (where rapid evolution of PRDM9 would quickly erode the signal of introgression), although further investigation is needed to test this hypothesis – ideally, as the number of high-quality archaic genomes continues to increase in the next years. Concretely, these results show that 1) iSMC can be used to extract information from archaic genomes; 2) hotspot turn-over driven by PRDM9 not only is responsible for short timescale differences in the location of cross-over events, but the accumulation of such differences is fast enough to have already impacted the landscape of *historical* recombination along the human lineage; and 3) in hominids, the spatial distribution of cross-over events changes at a reasonably constant rate, indicating that it evolves – to a large extent – neutrally.

**Figure 4.**
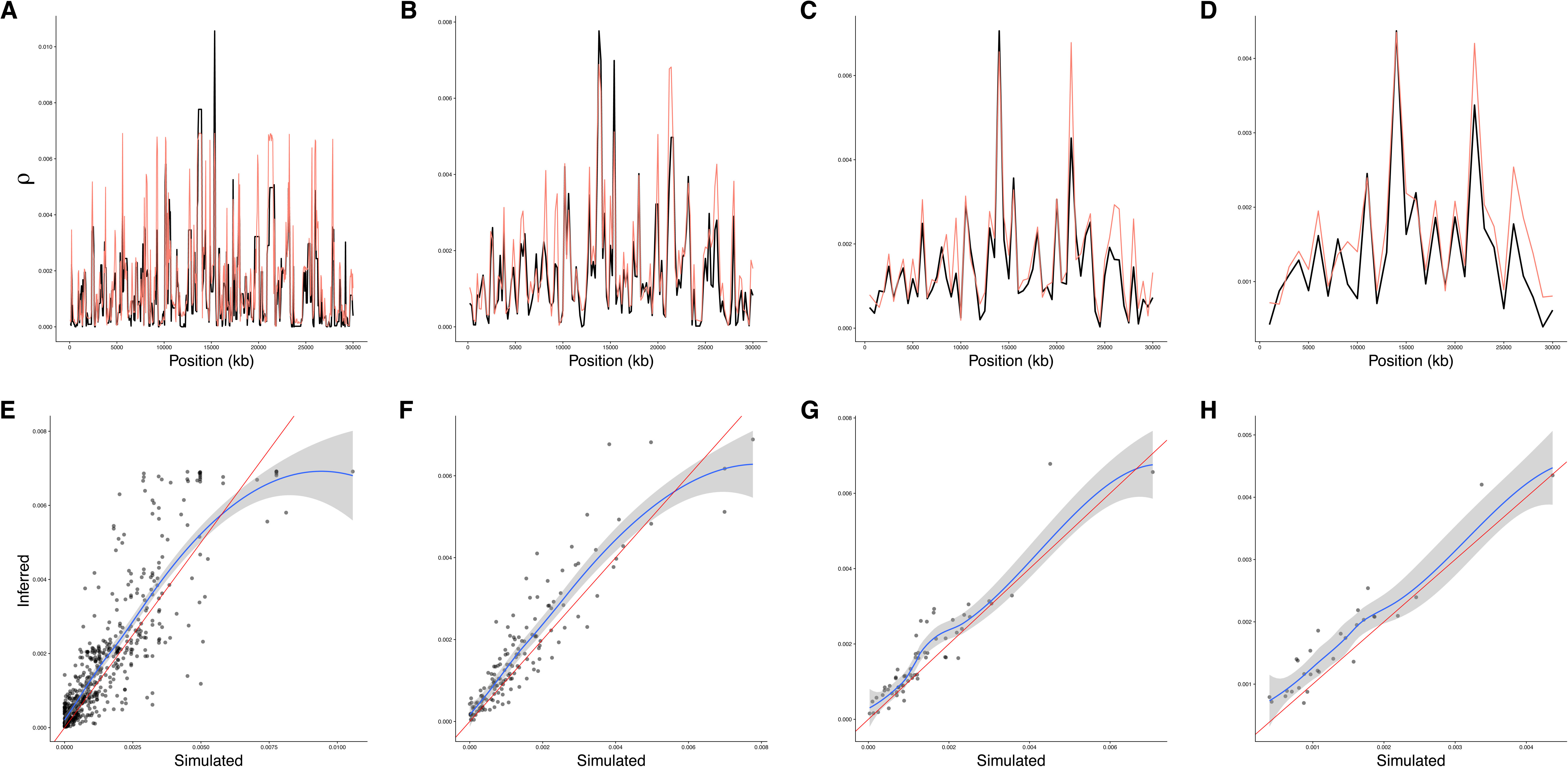
**Evolution of the fine-scale recombination landscape in hominids. A**, Dendogram based on pair-wise correlations between whole-genome recombination maps at the 50 kb scale. **B**, principal component analysis of the same data.

## DISCUSSION

Our analyses have shown that iSMC is able to infer accurate recombination maps from single pairs of genomes. Nevertheless, the correlations between statistical and experimental maps for the three species are consistently lower than the correlations obtained during simulations. While this difference can be partly explained by technical noise (e.g., sequencing or SNP calling errors), there are alternative – and conceptually engaging – explanations for it. First, the different types of data used by experimental and statistical methods imply that they measure different facets of recombination. While experimental maps reflect the average cross-over rate (*r*) between markers (i.e., a snapshot of the landscape at present-day generation), the population rate ρ (4.Ne.*r*) estimated by iSMC depends on ancestral population sizes as well as on the variation of Ne along the genome, and reflects the time-average of the historical recombination rate *r*. Since regions of high *r* tend to erode quickly^64,65^, the evolution of the recombination landscape over time implies that ρ may be quite different from *r* (and more meaningful than *r* in the context of population genomics^57^). Second, biological processes that affect Ne locally (therefore distorting the distribution of genealogies) but are unaccounted for by our model will affect LD without reflecting the recombination rate. Among these, introgression and selective sweeps (through the rapid increase of the frequency of advantageous haplotypes) can introduce a substantial bias if prevalent along the genome. In a similar vein, background selection distorts the distribution of genealogies towards lower TMRCAs^72^, and should be common near regions under purifying selection such as coding regions.

In order to study processes such as population structure^62^ and speciation^57,73^, population genomics generates high quality whole-genome sequences, with a small number of individuals representing each of several sub-populations^73–76^. Due to the importance of linkage in ancestral inference, recombination maps are key to analysing these data – and because of its power with restricted sample sizes, iSMC is well suited for the task. We have demonstrated its accuracy in species with contrasting levels of diversity, demographic histories and selective pressures, and posit that it will be useful for investigation in other species. Not only will such maps aid the interpretation of diversity in non-model organisms, but a picture of the recombination landscape in different groups will tell us about the nature of recombination itself^77^. Open questions include whether the recombination landscape is associated with large-scale genome architecture and how variation in the recombination landscape relates to life history and ecological traits. Finally, as ancient DNA samples become more common (including species other than humans^78^), it will be possible to obtain maps from extinct *taxa*, granting the opportunity to study the evolution of the recombination landscape with unprecedented resolution^45,79^.

## METHODS

### The Markov-modulated Hidden Markov Model framework

SMC models discretise a distribution of coalescence times into *t* intervals to implement a discrete space Hidden Markov Model (HMM) with *t* x *t* transition matrix:

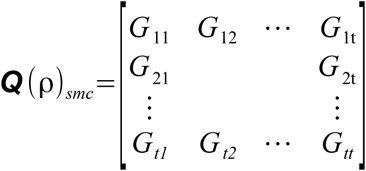

where *G*_ij_ (the transition probabilities between genealogies *i* and *j*) is a function of ancestral coalescence rates and the global parameter ρ, which is assumed to be constant along the genome^40,44^. The key innovation in iSMC is to relieve this assumption by letting ρ vary along the genome, following its own Markov process, where values drawn from an a priori distribution are used to compute the transition probabilities between genealogies. Let *R* be any strictly positive probability distribution with mean = 1.0 describing the variation in recombination along the genome. If *R* is discretised into *k* categories of equal density, the possible values that ρ can *r*_j_ * ρ_0_, where *r*_j_ is the *j*th *R* category and ρ_0_ is the genome-wide average recombination rate. Our Markov model (inspired by the observation that the distribution of cross-over events is not random, but clustered in regions of similar values) states that the probability distribution of *R* at position x + 1 only depends on the the distribution at position x. We consider the case where the transition probability between any two *R* categories (*P*_ij_) is identical and equivalent to one auto-correlation parameter (δ). The transition matrix of this Markovian process is simply:

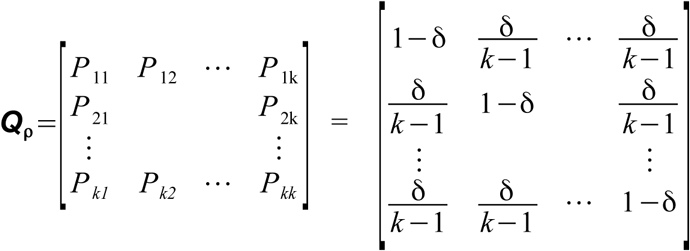

Because ρ is a parameter of the SMC, variation in the recombination rate affects the transition probabilities between genealogies (**Q_SMC_**). Since spatial variation in ρ is modeled as a Markovian process, the combined process is said to be Markov-modulated by ρ, leading to a Markov-modulated HMM. If *t* is the number of discrete genealogies of the SMC, and *k* is the number of discretised ρ categories, then the Markov-modulated HMM is a HMM with *n* = *t* x *k* hidden states (Figure 1). We further disallow transitions between hidden states where both the genealogies and ρ values change simultaneously. In this case, the transition matrix of the Markov-modulated process, **Q_iSMC_**, is given by:

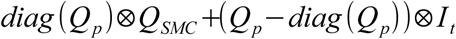

where û denotes the Kronecker product and **I_t_** the identity matrix of dimension *t*. Since many of its elements are reduced to zero, the matrix can be transversed efficiently:

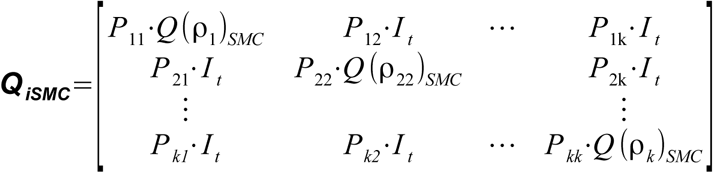

In other words, **Q_iSMC_** is a composition of *k*^2^ square sub-matrices of dimension *t*. The main diagonal sub-matrices are **Q_SMC_** assembled using the corresponding ρ value for category *k* (and scaled by 1 – δ), while the off-diagonal sub-matrices are identity matrices which have δ as main diagonal elements.

### Modelling spatial variation in recombination rates

iSMC implements three models of spatial variation in the recombination rate. We first use a Gamma probability density function with a single parameter (α = beta), which constrains it to have a mean equal to 1.0. After discretisation into *k* categories of equal density, the mean value inside each category is drawn to scale the genome-wide average ρ during integration over all recombination rates in the forward recursion (**equation 1**). In our simulation study, since we used a continuous Gamma distribution to draw values of the recombination landscape, we used this model to infer recombination maps. In the second model, we extend the gamma distribution by adding a category that represents the intensity of the recombination rate in sharp hotspots (parameter *H*). Since hotspots are narrow relative to the extension of the background recombination rate, we use extra parameters to accommodate this effect. As before, δ is the transition probability between gamma categories, and we introduce *u* as the transition probability from any gamma category to *H*, and *v* as the transition probability from *H* to any gamma category. The third model is obtained by letting the number of discretised categories of the Gamma distribution equal to 1, such that it becomes a probability mass function of two categories (**Supplemental Note**).

### Model selection and computation of the posterior recombination landscape

iSMC works in two steps: (1) fitting models of recombination rate variation and (2) inferring recombination maps based on the selected model. During step 1, the model parameters are optimized by maximizing the likelihood using the Powell multi-dimensions procedure^80^, which is computed for the entire sequence by applying the forward recursion of the HMM^48^ at every position *x* of the alignment:

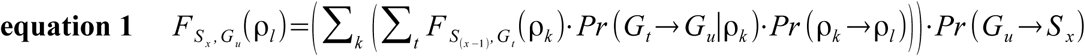

where we integrate over all *k* discretized values of ρ and over all *t* TMRCA intervals. The transition between genealogies (*G_t_*→*G_u_*) is a function of both the focal recombination rate (ρ_k_) and the ancestral coalescence rates, and *G_u_* →*S_x_* represents the emission probability from *G_u_* to the observed binary state at position *x* (**Supplemental Note**). In case AIC favours one of the heterogeneous models, in step 2 iSMC uses the estimated parameters to estimate the posterior average ρ for all sites in the genome. To this end, it first uses the so-called posterior decoding method^48^ as implemented in zipHMMlib^81^ to compute the posterior probability of every hidden state at each position in the sequence. Since in the Markov-modulated HMM the hidden states are pairs of ρ categories and TMRCA intervals, this results in joint probability distributions of recombination values and coalescence times, for all sites in the genome. Thus, prior to computing the posteror average, iSMC obtains the marginal distribution over recombination rates: if ^n^**Y**_x_ is the *n*-dimensional vector containing posterior probabilities for all *n* hidden states at position *x*, where *n* is the product of *t* TMRCA intervals and *k* ρ categories, iSMC obtains the *k*-dimensional vector ^k^**Z**_x_ by summing, for all *k*, the posterior probabilities of every hidden state *n* matching ρ category *k*. Finally, iSMC uses ^k^**Z**_x_ to scale the corresponding values of the discretized distribution of ρ, such that the posterior average ρ for position *x* is 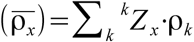.

### Modelling complex demographic histories

The original HMM implementation of the SMC^43^ uses the expectation-maximisation algorithm to optimise transition probabilities, where the actual targets of inference – the coalescence rates at each time interval – are hyper-parameters of the model. Here we use cubic spline interpolation^42^ to map coalescence rates at time boundaries, which are then assumed to be piecewise constant for the duration of each interval. Because we use three internal splines knots (i.e., the demographic history is divided into four epochs wherein a cubic curve is fitted), the number of parameters is substantially reduced in our model – in particular when a fine discretisation of TMRCA is employed.

### Simulation study

*Four scenarios of spatial variation in ρ*. We simulated a piecewise constant recombination rate along the genome by drawing values from a gamma distribution with parameters α and β, and segment lengths from a geometric distribution with mean length g. We considered four possible scenarios where α = β = 0.5 or 5.0, and g = 100 kb or 1 Mb. For each of the four combinations, we simulated 10 independent pairs of two 30 Mb haploid chromosomes under a constant population size model, assuming θ = 0.003 and ρ = 0.0012. For each of the following simulated scenarios, we focus on the landscape with α = 0.5 and g = 100 kb. All scenarios share base parameters values (θ, ρ, sequence length and sample size).

*Demographic history.* We simulated two demographic scenarios. First, a 20-fold population expansion 0.01 coalescent time units ago; second, a 20-fold population bottleneck 0.5 coalescent time units ago. Translating these coalescent times based on effective population sizes, the expansion happened 600 generations ago, and the bottleneck happened 30,000 generations ago.

*Introgression events.* We simulated two introgression scenarios where a source population introduces a pulse of genetic material into a target population. In both scenarios, the split between source and target populations happened 2.0 coalescent time units ago, and the source replaces 10% of the genetic pool of the target. In the first scenario, secondary contact happened 0.125 coalescent time units ago; in the second, it happened 0.25 coalescent time units ago. Translating these coalescent times based on effective population sizes, the split between population happened 120,000 generations ago, and the introgression events happened 7,500 and 15,000 generations ago, respectively.

*Variation in the mutation rate.* We simulated a piecewise constant mutation rate along the genome by drawing rate values from a uniform distribution and segment lengths from a geometric distribution with mean length f, where f is either 20 kb or 500 kb. The uniform distribution generating scaling factors of θ has mean = 5.05 instead of 1.0. In this case, the expected genome-wide average θ = 0.015. The reason for that is our focus on the spatial distribution of θ itself. If the landscape had mean = 0.003, its highly heterogeneous nature would scale θ down to values well below ρ (0.0012) too often along the 30 Mb sequence. The ensuing loss of signal (due to low SNP density) would result in poorly inferred maps that display low correlations with the simulations, not because of spatial *heterogeneity* in θ (local transitions), but instead because the ratio θ / ρ would be too low in many windows across the chromosome.

### Code availability

Source code for iSMC, as well as executable files with an example dataset, can be obtained at [github repository will be made public upon publication]. R scripts used both for simulations and data analyses are available in **Supplemental Data 1**.

### Data analysis

Model selection followed by inference of recombination maps in the three species studied (Table 1) was performed using publicly available sequences (chromosome 2L from haploid pairs ZI161 / ZI170, ZI179 / ZI191 and ZI129 / ZI138 in the Drosophila Population Genomics Project Phase 3^82^; chromosome 1 from haploid pairs Zt09 / Zt150, Zt154 / Zt155 and Zt05 / Zt07 for *Zymoseptoria tritici*^31^; chromosome 10 from three Finnish individuals (LP6005442-DNA_C10, LP6005442-DNA_D10, LP6005592-DNA_A02) available in the Simons Genome Diversity Project^62^ for humans). In the first two species, gaps and unknown nucleotides in the sequences (in FASTA format) were assigned as missing data, whereas in Humans the available strict mask for the dataset was applied after parsing the VCF files.

iSMC was fitted four times to each pair of genomes of the three species: 1) with 40 discretised time intervals and a model of variation in ρ based on a Gamma distribution with five discretised categories; 2) with 40 discretised time intervals and a model of variation in ρ based on an extended Gamma distribution with four discretised categories and an additional “Hotspot” category; 3) with 40 discretised time intervals and a model of variation in ρ based on a probability mass function of two categories; 4) with 100 discretised time intervals and a model of variation in ρ based on a probability mass function of two categories. In each case, Kendall correlation of ranks was computed between the resulting recombination landscape and available genetic maps both at the fine scale (100 kb for *Drosophila*, 20 kb for *Zymoseptoria tritici* and 50 kb for Humans) and at the large scale (1 Mb for *Drosophila*, 200 kb for *Zymoseptoria tritici* and 1 Mb for Humans).

Whole-genome sequence data were used for in-depth analyses of the recombination landscape in the hominid clade. Model fit (based on the “Hotspot” distribution) and inference of recombination maps was performed independently on each chromosome. The individual IDs within the Simons Genome Diversity Project^62^ and corresponding population of origin of the nine contemporary modern humans are as follows: African (Yoruban): LP6005442-DNA_A02, LP6005442-DNA_B02 and SS6004475; Asian (Dai Chinese): LP6005441-DNA_D04, LP6005443-DNA_B01 and LP6005592-DNA_D03; European (Finnish): LP6005442-DNA_C10, LP6005442-DNA_D10, LP6005592-DNA_A02. The available strict mask for the dataset was applied to assign low-quality positions as missing data. The four ancient DNA samples were downloaded from the server at the Max Planck Institute for Evolutionary Anthropology in Leipzig (http://cdna.eva.mpg.de/neandertal/Vindija/VCF) in May 2018. Since these are complete VCF files where all callable positions are reported, no mask was used and absent positions were assigned as missing data.

The analyses of modern and archaic datasets was performed considering only positions present in the DECODE genetic map. The dendogram presented in Figure 4 was obtained by hierarchical clustering (using complete linkage) of pair-wise distances computed from 1 – τ, where τ is the Kendall correlation of ranks between two individual recombination maps. Recombination maps for the entire archaic sequences are available as a resource in **Supplemental Data 2**.

## Acknowledgements

The authors thank Alice Feurtey, Asger Hobolth, Bernhard Haubold, Eva Stukenbrock, Fabian Klötzl, Kai Zeng, Pier Palamara and Stephan Schiffels for fruitful discussions about this work. JYD acknowledges funding from the Max Planck Society. This work was supported by a grant from the German Research Foundation (Deutsche Forschungsgemeinschaft) attributed to JYD, within the priority program (SPP) 1590 “probabilistic structures in evolution”.

## Author Contributions

JYD and GVB conceived and designed the study, analysed the datasets and wrote the manuscript. GVB developed the software. GVB and NP performed the simulation study.

The authors declare that there are no conflict of interest regarding the publication of this article.

## SUPPLEMENTAL FIGURE LEGENDS

**Figure S1. Inference of recombination maps in the presence of a heterogeneous mutation landscape.** Points are coloured as a function of the local ratio θ / ρ (>= 1 green, < 1 pink). Magenta dashed line represents y = x. **A**, both ρ and θ are binned into 50 kb windows. Solid lines represent loess smoothing curves fitted to each class ofθ / ρ, with 95% confidence intervals aroung them (shaded). **B**, both ρ and θ were binned into 500 kb windows.

**Figure S2. Evolution of the large-scale recombination landscape in hominids. A-C**, Dendogram based on pair-wise correlations between whole-genome recombination maps at the 50 kb, 200 kb and 1 Mb scales. **D-F**, principal component analysis of the same data.

## REFERENCES

1. Keightley, P. D. & Otto, S. P. Interference among deleterious mutations favours sex and recombination in finite populations. Nature 443, 89–92 (2006).

2. Hill, W. G. & Robertson, A. The effect of linkage on limits to artificial selection. Genet. Res. 8, 269–294 (1966).

3. Smith, J. M. & Haigh, J. The hitch-hiking effect of a favourable gene. Genet. Res. 89, 391– 403 (2007).

4. Ellegren, H. & Galtier, N. Determinants of genetic diversity. Nat. Rev. Genet. 17, 422–433 (2016).

5. Boulton, a, Myers, R. S. & Redfield, R. J. The hotspot conversion paradox and the evolution of meiotic recombination. Proceedings of the National Academy of Sciences of the United States of America 94, 8058–8063 (1997).

6. Myers, S. et al. Drive against hotspot motifs in primates implicates the PRDM9 gene in meiotic recombination. Science 327, 876–879 (2010).

7. Kong, A. et al. Fine-scale recombination rate differences between sexes, populations and individuals. Nature 467, 1099–1103 (2010).

8. Kawakami, T. et al. Whole-genome patterns of linkage disequilibrium across flycatcher populations clarify the causes and consequences of fine-scale recombination rate variation in birds. Mol. Ecol. 26, 4158–4172 (2017).

9. Dumont, B. L. & Payseur, B. A. Genetic Analysis of Genome-Scale Recombination Rate Evolution in House Mice. PLOS Genetics 7, e1002116 (2011).

10. Baudat, F. et al. PRDM9 is a major determinant of meiotic recombination hotspots in humans and mice. Science 327, 836–840 (2010).

11. Auton, A. et al. A fine-scale chimpanzee genetic map from population sequencing. Science 336, 193–198 (2012).

12. Singhal, S. et al. Stable recombination hotspots in birds. Science 350, 928–932 (2015).

13. Heil, S., S, C., Ellison, C., Dubin, M. & Noor, M. A. F. Recombining without Hotspots: A Comprehensive Evolutionary Portrait of Recombination in Two Closely Related Species of Drosophila. Genome Biol Evol 7, 2829–2842 (2015).

14. Brand, C. L., Cattani, M. V., Kingan, S. B., Landeen, E. L. & Presgraves, D. C. Molecular Evolution at a Meiosis Gene Mediates Species Differences in the Rate and Patterning of Recombination. Current Biology 28, 1289–1295.e4 (2018).

15. Kohl, K. P., Jones, C. D. & Sekelsky, J. Evolution of an MCM complex in flies that promotes meiotic crossovers by blocking BLM helicase. Science 338, 1363–1365 (2012).

16. Cutter, A. D. & Payseur, B. A. Genomic signatures of selection at linked sites: unifying the disparity among species. Nature reviews. Genetics 14, 262–74 (2013).

17. Wang, J., Street, N. R., Scofield, D. G. & Ingvarsson, P. K. Natural Selection and Recombination Rate Variation Shape Nucleotide Polymorphism Across the Genomes of Three Related Populus Species. Genetics 202, 1185–1200 (2016).

18. Schumer, M. et al. Natural selection interacts with recombination to shape the evolution of hybrid genomes. Science eaar3684 (2018). doi:10.1126/science.aar3684

19. Murray, G. G. R. et al. Natural selection shaped the rise and fall of passenger pigeon genomic diversity. Science 358, 951–954 (2017).

20. Martin, S. H. & Jiggins, C. D. Interpreting the genomic landscape of introgression. Curr. Opin. Genet. Dev. 47, 69–74 (2017).

21. Visscher, P. M. et al. 10 Years of GWAS Discovery: Biology, Function, and Translation. The American Journal of Human Genetics 101, 5–22 (2017).

22. Duret, L. & Galtier, N. Biased gene conversion and the evolution of mammalian genomic landscapes. Annu Rev Genomics Hum Genet 10, 285–311 (2009).

23. Kostka, D., Hubisz, M. J., Siepel, A. & Pollard, K. S. The Role of GC-Biased Gene Conversion in Shaping the Fastest Evolving Regions of the Human Genome. Mol Biol Evol 29, 1047–1057 (2012).

24. Bolívar, P., Mugal, C. F., Nater, A. & Ellegren, H. Recombination Rate Variation Modulates Gene Sequence Evolution Mainly via GC-Biased Gene Conversion, Not Hill–Robertson Interference, in an Avian System. Molecular Biology and Evolution 33, 216–227 (2016).

25. Glémin, S. et al. Quantification of GC-biased gene conversion in the human genome. Genome Res. 25, 1215–1228 (2015).

26. Stumpf, M. P. H. & McVean, G. a. T. Estimating recombination rates from population-genetic data. Nature Reviews Genetics 4, 959–968 (2003).

27. Rosenberg, N. A. & Nordborg, M. Genealogical Trees, Coalescent Theory and the Analysis of Genetic Polymorphisms. Nature Reviews Genetics 3, 380–390 (2002).

28. McVean, G. A. T. et al. The fine-scale structure of recombination rate variation in the human genome. Science 304, 581–584 (2004).

29. Auton, A. & McVean, G. Recombination rate estimation in the presence of hotspots. Genome Res 17, 1219–1227 (2007).

30. Chan, A. H., Jenkins, P. A. & Song, Y. S. Genome-Wide Fine-Scale Recombination Rate Variation in Drosophila melanogaster. PLOS Genetics 8, e1003090 (2012).

31. Stukenbrock, E. H. & Dutheil, J. Y. Fine-Scale Recombination Maps of Fungal Plant Pathogens Reveal Dynamic Recombination Landscapes and Intragenic Hotspots. Genetics 208, 1209–1229 (2018).

32. Hudson, R. R. Two-Locus Sampling Distributions and Their Application. Genetics 159, 1805–1817 (2001).

33. Hudson, R. R. & Kaplan, N. L. Statistical properties of the number of recombination events in the history of a sample of DNA sequences. Genetics 111, 147–164 (1985).

34. Hudson, R. R. & Kaplan, N. L. The coalescent process in models with selection and recombination. Genetics 120, 831–840 (1988).

35. Wiuf, C. & Hein, J. Recombination as a point process along sequences. Theoretical population biology 55, 248–59 (1999).

36. Griffiths, R. C. & Marjoram, P. An ancestral recombination graph. in Progress in population genetics and human evolution 257–270 (Springer, 1997).

37. Griffiths, R. C. & Marjoram, P. Ancestral inference from samples of DNA sequences with recombination. J. Comput. Biol. 3, 479–502 (1996).

38. Hein, J., Schierup, M. & Wiuf, C. Gene Genealogies, Variation and Evolution: A primer in coalescent theory. (Oxford University Press, 2004).

39. McVean, G. A. T. & Cardin, N. J. Approximating the coalescent with recombination. Philosophical Transactions of the Royal Society B: Biological Sciences 360, 1387–1393 (2005).

40. Marjoram, P. & Wall, J. D. Fast ‘coalescent’ simulation. BMC Genetics 7, 16–16 (2006).

41. Wilton, P. R., Carmi, S. & Hobolth, A. The SMC Is a Highly Accurate Approximation to the Ancestral Recombination Graph. Genetics 200, 343–355 (2015).

42. Terhorst, J., Kamm, J. A. & Song, Y. S. Robust and scalable inference of population history from hundreds of unphased whole-genomes. Nat Genet 49, 303–309 (2017).

43. Li, H. & Durbin, R. Inference of human population history from individual whole-genome sequences. Nature 475, 493–496 (2011).

44. Schiffels, S. & Durbin, R. Inferring human population size and separation history from multiple genome sequences. Nature Genetics 46, 919–925 (2014).

45. Munch, K., Schierup, M. H. & Mailund, T. Unraveling recombination rate evolution using ancestral recombination maps. Bioessays 36, 892–900 (2014).

46. Munch, K., Mailund, T., Dutheil, J. Y. & Schierup, M. H. A fine-scale recombination map of the human-chimpanzee ancestor reveals faster change in humans than in chimpanzees and a strong impact of GC-biased gene conversion. Genome Res. 24, 467–474 (2014).

47. Dutheil, J. Y. Hidden Markov Models in Population Genomics. Methods Mol. Biol. 1552, 149–164 (2017).

48. Durbin, R., Eddy, S. R., Krogh, A. & Mitchison, G. Biological sequence analysis: Probabilistic models of proteins and nucleic acids. (Cambridge University Press, 1998). doi:10.1017/CBO9780511790492

49. Slatkin, M. Linkage disequilibrium--understanding the evolutionary past and mapping the medical future. Nature reviews. Genetics 9, 477–85 (2008).

50. McVean, G. A. T. A genealogical interpretation of linkage disequilibrium. Genetics 162, 987–991 (2002).

51. Wirtz, J., Rauscher, M. & Wiehe, T. Topological linkage disequilibrium calculated from coalescent genealogies. Theoretical Population Biology (2018). doi:10.1016/j.tpb.2018.09.001

52. Akaike, H. A new look at the statistical model identification. IEEE Transactions on Automatic Control 19, 716–723 (1974).

53. Staab, P. R., Zhu, S., Metzler, D. & Lunter, G. Scrm: Efficiently simulating long sequences using the approximated coalescent with recombination. Bioinformatics 31, 1680–1682 (2015).

54. Dapper, A. L. & Payseur, B. A. Effects of Demographic History on the Detection of Recombination Hotspots from Linkage Disequilibrium. Mol. Biol. Evol. 35, 335–353 (2018).

55. Johnston, H. R. & Cutler, D. J. Population demographic history can cause the appearance of recombination hotspots. Am. J. Hum. Genet. 90, 774–783 (2012).

56. Kamm, J. A., Spence, J. P., Chan, J. & Song, Y. S. Two-Locus Likelihoods Under Variable Population Size and Fine-Scale Recombination Rate Estimation. Genetics 203, 1381–1399 (2016).

57. Martin, S. H., Davey, J., Salazar, C. & Jiggins, C. Recombination rate variation shapes barriers to introgression across butterfly genomes. bioRxiv 297531 (2018). doi:10.1101/297531

58. Grossen, C., Keller, L., Biebach, I., Consortium, T. I. G. G. & Croll, D. Introgression from Domestic Goat Generated Variation at the Major Histocompatibility Complex of Alpine Ibex. PLOS Genetics 10, e1004438 (2014).

59. Francioli, L. C. et al. Genome-wide patterns and properties of *de novo* mutations in humans. Nature Genetics 47, 822–826 (2015).

60. Hartmann, F. E., McDonald, B. A. & Croll, D. Genome-wide evidence for divergent selection between populations of a major agricultural pathogen. Molecular Ecology 27, 2725–2741

61. Croll, D., Lendenmann, M. H., Stewart, E. & McDonald, B. A. The impact of recombination hotspots on genome evolution of a fungal plant pathogen. Genetics 201, 1213–1228 (2015).

62. Mallick, S. et al. The Simons Genome Diversity Project: 300 genomes from 142 diverse populations. Nature (2016). doi:10.1038/nature18964

63. Paigen, K. & Petkov, P. M. PRDM9 and Its Role in Genetic Recombination. Trends Genet. 34, 291–300 (2018).

64. Coop, G. & Myers, S. R. Live hot, die young: Transmission distortion in recombination hotspots. PLoS Genetics 3, 0377–0386 (2007).

65. Latrille, T., Duret, L. & Lartillot, N. The Red Queen model of recombination hot-spot evolution: a theoretical investigation. Phil. Trans. R. Soc. B 372, 20160463 (2017).

66. Lesecque, Y., Glémin, S., Lartillot, N., Mouchiroud, D. & Duret, L. The Red Queen Model of Recombination Hotspots Evolution in the Light of Archaic and Modern Human Genomes. PLOS Genetics 10, e1004790 (2014).

67. Slatkin, M. & Racimo, F. Ancient DNA and human history. PNAS 113, 6380–6387 (2016).

68. Prüfer, K. et al. The complete genome sequence of a Neanderthal from the Altai Mountains. Nature 505, 43–49 (2013).

69. Green, R. E. et al. A draft sequence of the Neandertal genome. Science (New York, N.Y.) 328, 710–22 (2010).

70. Meyer, M. et al. A High-Coverage Genome Sequence from an Archaic Denisovan Individual. Science 338, 222–226 (2012).

71. Fu, Q. et al. Genome sequence of a 45,000-year-old modern human from western Siberia. Nature 514, 445–449 (2014).

72. Palamara, P. F., Terhorst, J., Song, Y. S. & Price, A. L. High-throughput inference of pairwise coalescence times identifies signals of selection and enriched disease heritability. Nature Genetics 50, 1311–1317 (2018).

73. Brandvain, Y., Kenney, A. M., Flagel, L., Coop, G. & Sweigart, A. L. Speciation and Introgression between Mimulus nasutus and Mimulus guttatus. PLOS Genetics 10, e1004410 (2014).

74. Teng, H. et al. Population Genomics Reveals Speciation and Introgression between Brown Norway Rats and Their Sibling Species. Mol Biol Evol 34, 2214–2228 (2017).

75. Van Belleghem, S. M. et al. Patterns of Z chromosome divergence among Heliconius species highlight the importance of historical demography. Mol. Ecol. (2018). doi:10.1111/mec.14560

76. Delmore, K. E. et al. Comparative analysis examining patterns of genomic differentiation across multiple episodes of population divergence in birds. Evolution Letters 2, 76–87 (2018).

77. Stapley, J., Feulner, P. G. D., Johnston, S. E., Santure, A. W. & Smadja, C. M. Variation in recombination frequency and distribution across eukaryotes: patterns and processes. Phil. Trans. R. Soc. B 372, 20160455 (2017).

78. Librado, P. et al. Ancient genomic changes associated with domestication of the horse. Science 356, 442–445 (2017).

79. Moorjani, P. et al. A genetic method for dating ancient genomes provides a direct estimate of human generation interval in the last 45,000 years. PNAS 113, 5652–5657 (2016).

80. Guéguen, L. et al. Bio++: efficient extensible libraries and tools for computational molecular evolution. Mol. Biol. Evol. 30, 1745–1750 (2013).

81. Sand, A., Kristiansen, M., Pedersen, C. N. & Mailund, T. zipHMMlib: a highly optimised HMM library exploiting repetitions in the input to speed up the forward algorithm. BMC Bioinformatics 14, 339 (2013).

82. Lack, J. B. et al. The Drosophila Genome Nexus: A Population Genomic Resource of 623 Drosophila melanogaster Genomes, Including 197 from a Single Ancestral Range Population. Genetics 199, 1229–1241 (2015).

